# Soils-on-a-chip reveal unforeseen motility parameters of microconfined *Bradyrhizobium diazoefficiens*

**DOI:** 10.1101/2023.12.29.573673

**Authors:** Moniellen Pires Monteiro, Juan Pablo Carrillo, Nahuel Gutiérrez, Sofía Montagna, Aníbal R. Lodeiro, María Luisa Cordero, V. I. Marconi

**Affiliations:** Departamento de Física - Facultad de Ciencias Físicas y Matemáticas, Universidad de Chile, Santiago, Chile; Facultad de Matemática, Astronomía, Física y Computación, Universidad Nacional de Córdoba, Córdoba, Argentina; Instituto de Biotecnología y Biología Molecular - Facultad de Ciencias Exactas, Universidad Nacional de La Plata and CCT-La Plata CONICET, La Plata, Argentina; Laboratorio de genética - Facultad de Ciencias Agrarias y Forestales, Universidad Nacional de La Plata, La Plata, Argentina; IFEG-CONICET, Córdoba, Argentina

**Keywords:** soil-bacteria, soil-on-a-chip, microfluidics, biofertilizer, inoculant, sustainability, bioproducts, environment, soybean, legumes

## Abstract

Soil bacteria of the order of the Rhizobiales associate symbiotically with legume plants. Particulary, *Bradyrhizobium diazoefficiens* is a nitrogen-fixing symbiont of soybean, that helps to improve grain quality among other benefits. This bacterium possess two flagellar systems, which enable it to swim in water-saturated pores. However, the motility of B. diazoefficiens, which may be crucial for its competitiveness in root nodulation, has not been well understood. To address this knowledge gap, we designed and fabricated microfluidic soil-on-a-chip (SOC) devices that offer sustainable agriculture an original tool for directly visualizing bacterial behavior in confined-environments. Using these microdevices, we measured the population velocities and changes of direction along their paths for two strains of *B. diazoefficiens*, namely the wild-type and a mutant with only one flagellar system. Our detailed statistical analysis revealed that both strains exhibited reduced speeds and increased changes of direction of 180°, in channels of decreasing microscopic cross sectional area, down to a few microns. Interestingly, while the wild-type strain displayed faster swimming speeds in unconfined spaces, this advantage was negated in the SOCs that mimicked porous soils. Moreover, we employed the measured motility parameters to model and simulate *B. diazoefficiens* motion in SOC devices for extended periods and larger scales, enabling further predictions of diffusion in real soils. Thanks to miniaturization, microfabrication, and multidisciplinary knowledge, this study represents a significant breakthrough in soil bacteria field and methods, useful both for farmers and environment. Furthermore, the potential applications of this work extend to multiple beneficial bacteria widely used as biofertilizers.

---

Most bacterial species thriving in soil or aquatic environments display a flagella-driven self-propelled motion, hereafter referred to as flagellar motility, which enables them to explore the habitat, swimming in liquid media. Among these environmental microbial, the Rhizobiales order is of paramount importance because it contains most of the N_2_-fixing species symbiotic with legume plants. In this order, flagellar genes were detected in 141 complete genomes among 150 analysed so far (1.

Inoculation of legume crops with symbiotic rhizobia to profit from N_2_-fixation is routinely practiced since more than a century ago (2). Rhizobia are inoculated on seeds or sowing furrows from which they must access the infection targets at the root surfaces to invade the plant tissues and form N_2_-fixing nodules (2; 3). Therefore, rhizobia should move in the soil from the inoculation sites to the infection targets either passively dragged by water movement, or actively by flagellar motility. How this transit from the inoculation site to the infection targets is achieved is under debate since more than 50 years (4), and has not been studied through direct observation yet, according to our knowledge. Since rhizobial flagellar motion in the soil pores may be determinant for nodules occupation and crop yield (5), its understanding can represent major advances for the field of bioproducts, helping to achive a rational design of improved strains and inoculant application technologies (4; 6).

Most experiments aimed at studying rhizobial flagellar motility were carried out in homogeneous liquid environments, which are too reductionist with respect to the real situation in the soil. Unlike homogeneous liquids, the soil is a heterogeneous matrix filled by different sized pores and channels that are narrow and tortuous, wherein rhizobia must swim. In addition, the water content is not constant in soil, varying from flooding to several degrees of water deficit both in time and space. On the other hand, experiments carried out in soils or porous settings such as pots filled with inert substrates like perlite, vermiculite, sand, or their mixtures, indicated that rhizobial motility in soils is restricted to the soil flooded state, and its range is very limited in comparison to homogeneous liquids (4). However, these conclusions come from indirect observations, mostly rescuing bacteria trapped by roots, or evaluating competition for root colonization or nodulation between wild-type strains and derived mutants affected in flagellar or chemotaxis-related genes. Thus, there is a difficulty to integrate both lines of evidence, because many important details of rhizobial motility, such as swimming speed, changes of direction or types of trajectories, can be addressed only by direct microscopic observation of living swimming bacteria, which up to now, cannot be performed in soil samples. This important current issue inspired our multidisciplinary work on soil bacteria.

*Bradyrhizobium diazoefficiens* is the N_2_-fixing symbiont of soybean, the most important grain legume cropped worldwide. This bacterium has the unusual property of possessing two independent flagellar systems: a thick and short subpolar flagellum and thinner, longer lateral flagella. According to specific nutritional conditions, these two systems may be expressed simultaneously in liquid medium (7; 8). In addition, instability induced in the subpolar flagellum by inactivating the FliL_S_ protein induced a higher expression of the lateral flagella, suggesting the existence of a crosstalk between both systems (9).

Previously, we proposed the hypothesis that displacement of *B. diazoefficiens* in soil pores may be facilitated by flagellar swimming near soil pore walls (10). For this displacement, a trade-off between flagellar swimming and adhesion to pore walls must be established, on which lateral flagella may play a role (10). However, at the time we had no means of testing this hypothesis with direct observations of swimming within closed channels. Developments in microfluidics, that open doors to microbiology (11; 12; 13) and its applications (14; 15), has allowed us to prepare transparent microscopic chambers containing channels resembling the soil grain complex structure, and directly observe bacteria swimming within channels of different widths surrounding grains of different sizes (Fig. 1). Using these tools we were able to directly observe, quantify and compare the swimming of the *B. diazoefficiens* wild type (WT) and mutants devoid of subpolar (Δ*fliC*) or lateral (Δ*lafA*) flagellar filaments within artificial soils with different channel widths. These experiments in soil-on-a-chip (SOC) devices allowed us to perform a pioneer and detailed study of the flagellar swimming motility trajectories under microconfinement in conditions alike a flooded soil. Such contributions hope to offer new tools for direct visualization of soil bacteria that will be useful for the development of both new strains with optimal behavior, and innovative biotechnological applications related to biofertilizers inoculation methods, both key elements for the development of a future sustainable agriculture (16; 17; 3).

**Fig. 1.**
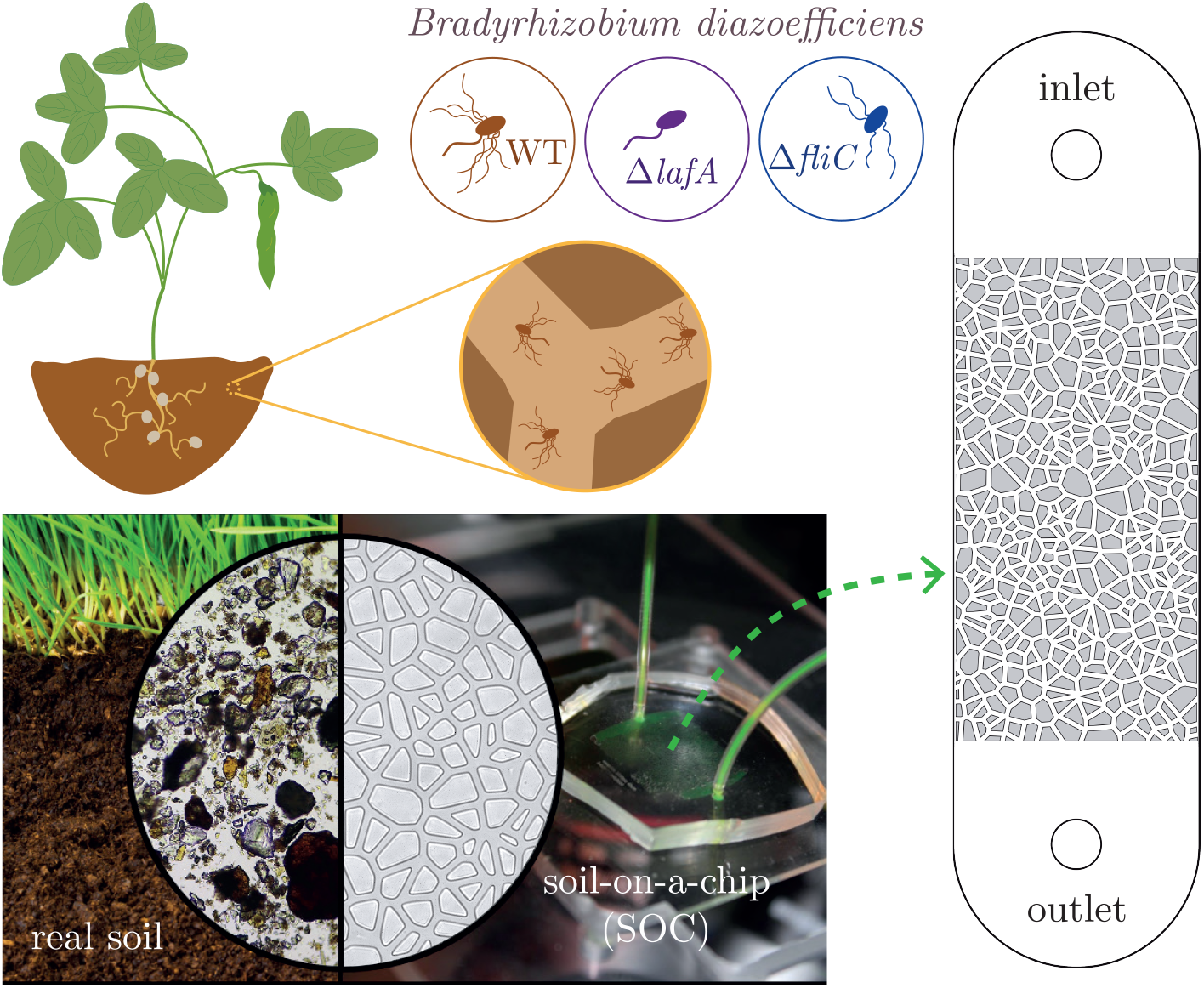
Direct observation of soil bacteria into miniaturized artificial soils. Flagellated soil bacteria navigate the soil by swimming through flooded pores between soil particles and grains. To visualize its swimming through a complex geometry with different degrees of confinement, an artificial soil model was used, both experimental and in-silico. Microfluidic soil-on-a-chip devices, as sketched in the zoom, were fabricated based on Voronoi tessellations (see Materials and Methods), with channels widths of increasing confinement (20 μm, 10 μm and 5 μm). Three different strains of *Bradyrhizobium diazoefficiens* were inoculated in the devices, a wild type, WT, with two flagellar systems and two mutants, one lacking the lateral flagellar system, Δ*lafA*, and the other lacking the subpolar one, Δ*fliC*, depicted as light-brown, violet, and blue bacteria, respectively.

## Reduction of swimming speed in artificial soils

We developed SOC devices to mimic the intricate geometry of flooded porous soil. Real soils are complex, highly heterogeneous structures, formed by grains whose size can range from micrometers to a few millimeters. In our devices, we fabricated grains of sizes between 5 μm and 100 μm, which represent silty and fine sandy soils. As grains reduce in size, so do the pores between the grains. Thus, our devices had different degrees of confinement, which was quantified by the width *w*_*i*_ of the channels where bacteria navigate (see Methods). Three different devices were fabricated with increasingly narrower channels, *w*_1_ = 20 μm, *w*_2_ = 10 μm, and *w*_3_ = 5 μm. In this way, we increment the degree of confinement by progressively halving the width of the channels. As a reference, the typical length of a *B. diazoefficiens* cell is about 1 μm or 2 μm when only considering the cell body, and about 10 μm when considering the complete cell including its flagella. Hence, the effects of walls is expected to increase as confinement becomes tighter, for example modifying the individual bacteria swimming strategies and the collective population transport properties.

Using bright field microscopy and digital imaging, we recorded and tracked the trajectory of thousands of bacteria of the three mutants (see Methods), swimming in the SOC devices. We observed that the vast majority of Δ*fliC* mutants (strain with lateral flagella only) moved irregularly, as trembling in a given position, without noticeable displacement (see supplementary videos), whereby we decided not to continue with the analysis of this mutant.

Figure 2 shows the probability density function (PDF) of the bacteria speed for the WT and Δ*lafA*, strains with both flagellar systems and with a subpolar flagellum only, respectively, swimming in the three devices. All experiments were repeated with at least three biological replicas, in order to assess data reproducibility and to obtain good statistics. In each experiment, bacteria trajectories were also measured in the inlet compartments (see Fig. 1), where bacteria swim in a quasi-two-dimensional chamber without grains or obstacles. The speed distributions obtained for each strain in these inlet regions are shown in the first row of Fig. 2 as a reference.

**Fig. 2.**
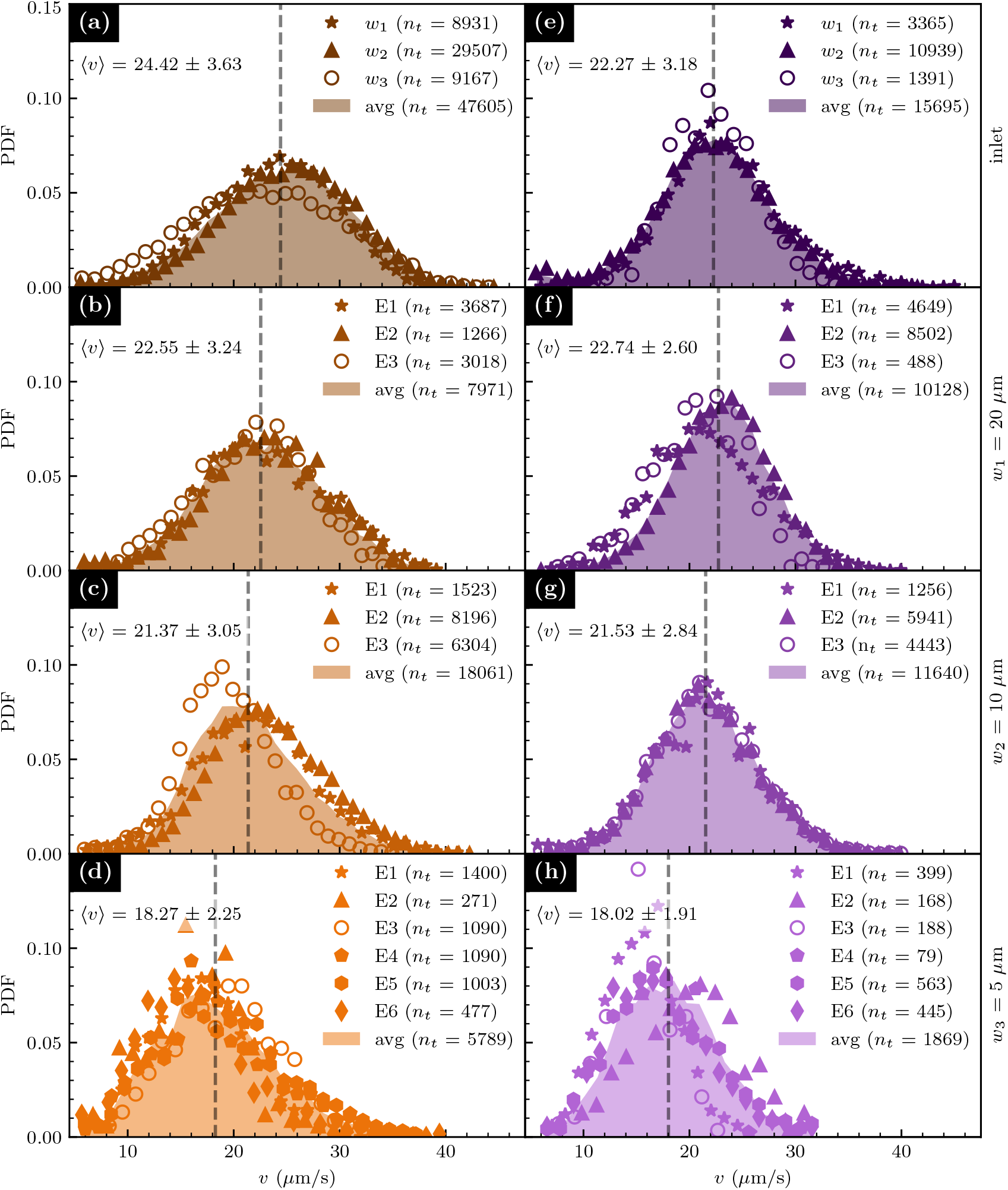
Speed distributions of bacteria in SOC. Speed probability density function (PDF) of WT [(a)–(d)] and Δ*lafA* [(e)–(h)] *B. diazoefficiens* strains are shown for measurements into: the inoculation site [panels (a) and (e)], a porous geometry with channels of width *w*_1_ = 20 μm [(b) and (f)], *w*_2_ = 10 μm [(c) and (g)], and *w*_3_ = 5 μm [(d) and (h)]. Symbols represent experimental distributions from 3 to 6 biological replicas (E_*i*_, with *i* = 1, …, 6), with a number of observed tracks *n*_*t*_ indicated for each case in the legends. Solid light colored backgrounds represent the average of all experiments (avg). Mean speed values *±* error (in units of μm*/*s) are given in all panels, and indicated with a dashed vertical line.

In general, differences in the speed PDFs between biological replicas are minor, and reflect some diversity among different cultures. However, when all data are combined, the resultant speed PDF for each population are smooth, reflecting the high quality statistics; they were obtained with thousands of cell tracks, each one lasting more than 0.5 s. This time of tracking is difficult to achieve in these kind of setups, and is roughly equivalent to a displacement of 12-15 body cells on average, a time-lapse during which bacteria swim in focus. In the inlets, the speed PDFs for both strains are single peaked and symmetric, with most of the cells swimming at speeds close to the maximum of the distribution and a small population of bacteria swimming at speeds smaller than 10 μm*/*s and larger than 35 μm*/*s. As the width of the channels decreases, the observed behavior is similar for both strains: the speed distributions remain single peaked, but the swimming speeds become progressively slower, which is manifested by a systematic shifting of the distributions toward the left. In the case of the WT, this shifting is already evident in the channels with width 20 μm, and continues shifting towards smaller speeds as the channel width decreases. In the case of the Δ*lafA*, the speed distribution in the widest channels does not differ appreciably with respect to the inlet. However, a systematic decrease in mean speed can be seen in the networks with channel width 10 μm and 5 μm.

We compare the speed PDFs of both strains in the different geometries in Fig. 3(a)–(d). In the inoculation chambers (Fig. 3(a)), the speed distributions of the WT and the mutant strain are appreciably different. The one for the WT is displaced to higher values compared to the speed distribution of the mutant. In quantitative terms, the mean speed of the WT in the inlet is (24.42±3.63) μm*/*s, and that of the single-flagellated Δ*lafA* is (22.27 ± 3.18) μm*/*s. That corresponds to a mean speed difference of almost 10 %. This behavior indicates that, on average, the presence of the two flagellar systems propels the WT bacteria with higher speed. While in the Δ*fliC* mutant the lateral flagella produce irregular and poor swimming, a complex cooperative behavior between the two flagellar systems appears to be at work in the case of the WT, allowing them to swim faster than the Δ*lafA* mutant. The speed distribution is also wider for the WT than for the Δ*lafA* strain in the inlet. This could be explained by natural variations in the number and length of lateral flagella, which causes population diversity in the case of the WT. In comparison, the Δ*lafA* mutant, only possessing a single flagellar system, has fewer degrees of freedom associated with the flagellar motor system, thus resulting in a more homogeneous population and a more concentrated speed distribution.

**Fig. 3.**
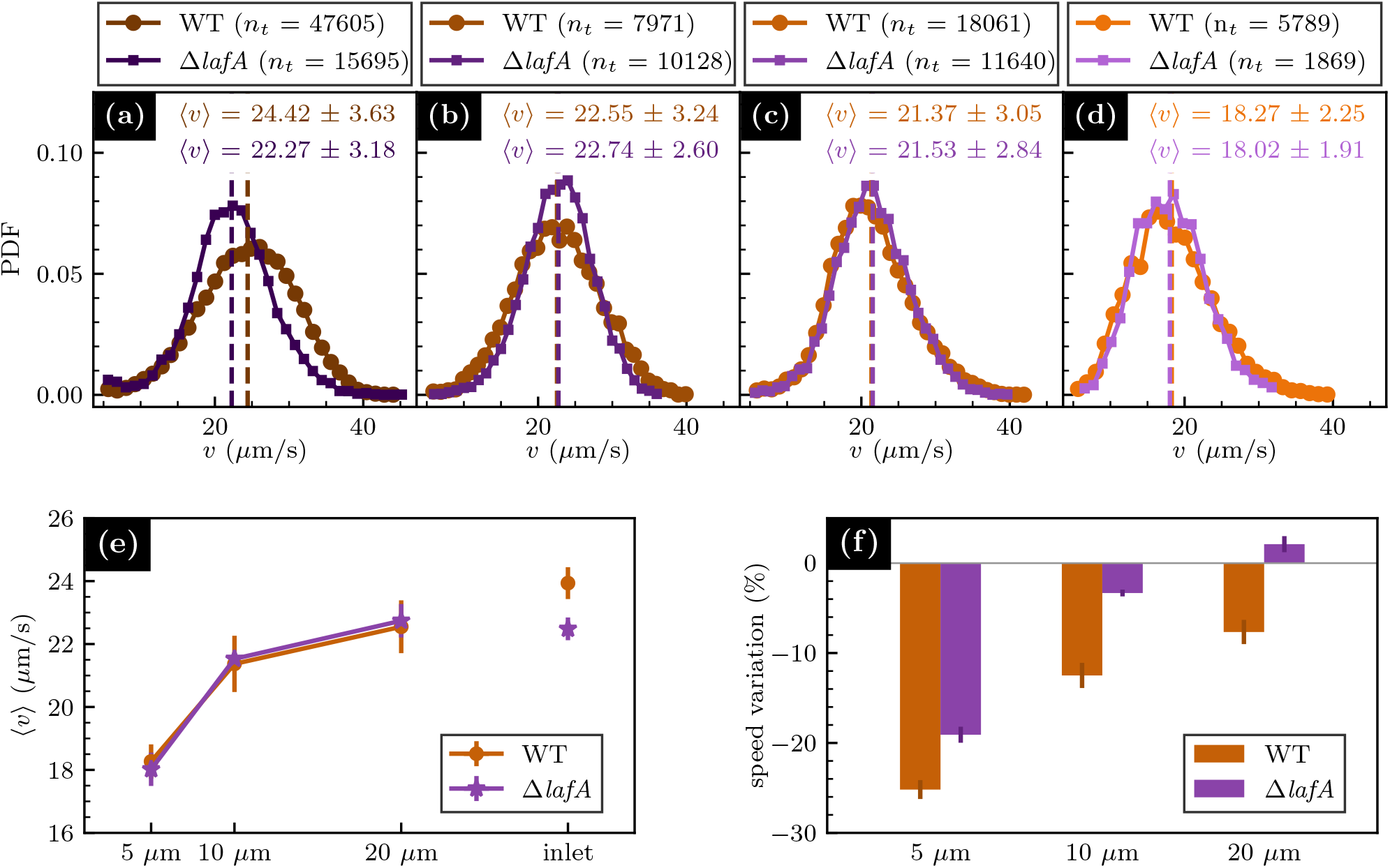
Average speed and speed variation ratio in artificial soils with different degrees of confinement. Speed PDF for WT and Δ*lafA* mutant in the inoculation chamber (a) and in artificial soil devices with different degrees of confinement: *w*_1_ = 20 μm (b), *w*_2_ = 10 μm (c), and *w*_3_ = 5 μm (d). (e) Bacteria average speed in the artificial soils as a function of the channel width. (f) SVR for each strain as a function of the channel width (see Eq. (1)). Vertical bars in both cases indicate the data corresponding errors.

As the available space to swim is reduced, the mean speed of both strains decreases, as explained previously. In the SOC devices, the average swimming speed coincide almost exactly for both strains in all channel widths including the widest one, even though on average the WT strain swims faster than the mutant in unconfined space (Fig. 3(b)–(c)). Moreover, as confinement increases, the speed distributions of both strains become more alike. In the channels of width 20 μm, the speed distribution of the WT is still wider than that of the Δ*lafA*, but in the channels of width 10 μm and 5 μm, the speed PDFs of both strains are almost identical. The only notable difference between both strains is that the speed distributions for the WT in the SOCs become asymmetric, systematically showing longer tails at large speeds than the Δ*lafA*. This reveals that, although the average speed of the overall population decreases in both strains, and the population diversity for the WT is reduced, a small but non negligible fraction of WT bacteria maintain high swimming speeds, even in the most confined geometries. On the contrary, the speed PDFs for the Δ*lafA* remain symmetric, showing that the overall population of bacteria decrease their swimming speed.

The average speed for each channel width is plotted in Fig. 3(e). The fall of mean speed is evident for the artificial soil with narrower channels. To quantify this speed reduction with respect to the mean speed in the inlet chamber, we define the speed variation ratio (SVR) as

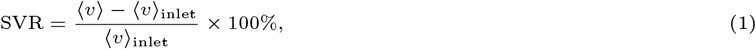

where ⟨*v*⟩_inlet_ is the average speed in the inlet for each strain. The SVR is presented in Fig. 3(f). In the widest channels, the SVR is about −7.5 % for the WT and 2 % for the mutant. As the confinement increases, the SVR falls for both strains, reaching −12.5 % and −25 % for the WT in the channels with width 10 μm and 5 μm, respectively. The corresponding SVR for the Δ*lafA* strain in the same confinements are −3 % and −19 %. In this case, the speed reductions are lower than in the WT, because the reference value in the inlet is lower for the mutant, but the average values in the artificial soils are very similar for both strains.

Overall, the results shown above indicate that confinement affects the swimming behavior of both strains. However, it appears to affect differently each strain, as can be seen when comparing the speed distributions of each strain to their respective distributions in the inlet. For the mutant inside the artificial soils, although the mean speed decreases systematically with decreasing channel width, the shape of the speed distribution remains almost unaltered; the height, width, and symmetry of the distribution remains approximately constant. For the WT strain, conversely, the speed distributions become narrower, taller, and asymmetric in the confining SOCs, and both in the mean speed and in the shape of the speed PDF, its behavior becomes similar to the Δ*lafA* mutant. The cooperative behavior between the two flagellar systems observed for WT swimming in the inlets appears to be lost, on average, when they swim in the artificial soils.

## Swimming strategies of B. diazoefficiens in porous media

An important component of the behavior of swimming bacteria is their specific strategies to rectify their direction of movement by reorienting during swimming. For soil bacteria, this behavior should be highly dependent on the specific porous media in which they navigate to live and survive. For this reason, we characterized the features of bacteria paths in all regions of the microdevices. Figure 4 shows different trajectory examples in the inlets and in the SOC for both the WT and the Δ*lafA* strains. These bacteria commonly reorient during their swimming. The angle Φ quantifies the change of direction (CHD) events. We define Φ as the positive difference in orientation angle between the final and initial direction of motion, as sketched in Fig. 4(a). When Φ is small, Φ ∈ [15°, 35°], the motion is considered highly persistent and bacteria move essentially forward. On the contrary, CHDs with Φ *>* 90° contribute to backward motion of the bacteria.

**Fig. 4.**
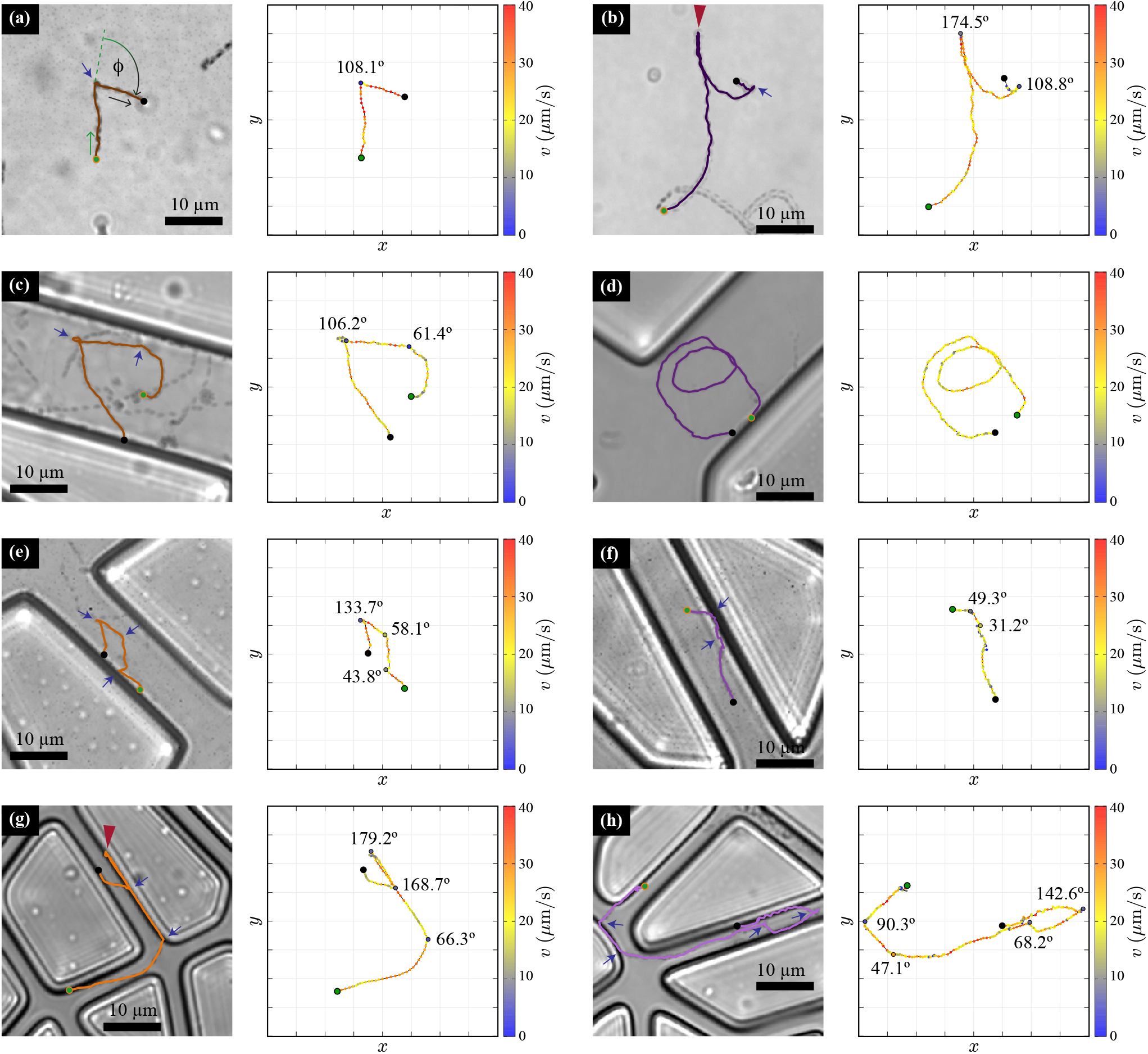
Swimming behavior of *B. diazoefficiens* in unconfined space and in SOC devices. Examples of trajectories followed by WT (left) and Δ*lafA* (right) at the unconfined inlets [(a) and (b)] and in the SOC microdevices with increasing confinement [*w*_1_ = 20 μm (c) and (d); *w*_2_ = 10 μm (e) and (f); and *w*_3_ = 5 μm (g) and (h)]. For each case, the trajectory is shown at the left and at the right the bacterium speed is presented along their path in color scale, with the angle values for each detected change of direction, Φ. Several snapshots are overlaid to show the trajectory of the bacteria for the time span of the tracks, and only one bacterium track is shown. The green and black dots mark, respectively, the start and end of the track. The blue arrows and red arrow heads mark changes of direction, (CHD) and run and reverses (RR), respectively. In (a) the reorientation event is marked with the definition of the reorientation angle, Φ. Videos of the trajectories are available in the Supplementary Material.

Figure. 4(a) presents a single, evident CHD for a WT bacterium, to show our definition of CHD and Φ. In this case, Φ = 108.1°. The track starts with the green dot and finishes at the black one. The blue arrow points at the reorientation event. The contiguous panel evidences the speed along the path, which fluctuates between 20 μm*/*s and 40 μm*/*s, with an important fall to less than 5 μm*/*s at the CHD.

One example of bacteria trajectory with CHDs for the Δ*lafA* strain in the inlet, along with the corresponding bacteria speed, is shown in Fig. 4(b). A remarkable behavior is the common existence of CHDs with an angle Φ ≈ 180°. These events occur in both strains and are reminiscent of the “Run and Reverse” (RR) behavior exhibited by some monotrichous bacteria under certain circumstances, such as *Vibrio alginolyticus* (18; 19) and *Caulobacter crescentus* (20; 21). Examples of RR events are marked with a red arrow head in Fig. 4. Close inspection of the videos suggests that RRs occur when the whole body of the bacterium turns in 180°, instead of reversing the swimming direction with the body in the same orientation (see videos in Supplementary Material). This implies that the reversal is not due to the flagella reversing their spinning direction. Due to noise, the RR angle is not necessarily equal to 180°. It is difficult to associate a threshold value for Φ to characterize a RR event, however, by inspection of the videos, we have decided to classify all CHDs with Φ *>* 160° as RRs.

In the channel networks, bacteria interact with bottom-top surfaces and the vertical boundaries. In the case with *w*_1_ = 20 μm, bacteria exhibit large trajectories and can perform more than one CHD in the channels before contacting the boundaries. Figure 4(c) shows one example trajectory of a WT bacterium swimming inside a channel and performing two CHDs before hitting the wall. In Fig. 4(d), one example trajectory of a Δ*lafA* cell can be observed. In this case, the swimmer circles twice between two close vertical walls without hitting them.

In the SOC with *w*_2_ = 10 μm, the interaction with boundaries becomes more common. Still, bacteria can swim in-plane for long trajectories and have CHDs before touching a vertical wall. Figures 4(e) and (f) show, respectively, trajectories of a WT and a Δ*lafA* in the SOC with width 10 μm. In both cases, the bacteria swim near the solid boundary for a short distance of approximately 5 μm before a CHD reorients them into the channel. We commonly observe this behavior, with in-plane path segments in which a bacterium swims close to the walls for a fraction of a second and then swims again into the channel, away from the wall.

In the network with channel width *w*_3_ = 5 μm, bacteria show qualitatively different behaviors. An example of a WT bacterium swimming in this device is presented in Fig. 4(g). The path presents a relatively long segment before a collision with a solid boundary. After hitting the wall, the bacterium aligns to it. Although the solid boundaries appear dark as well as bacteria, post-processing of the images allowed us to detect the cell swimming for a few seconds very near the wall, even after a RR. Eventually, another CHD causes the cell to cross the channel and align to the opposite wall. Similarly, the trajectory of a Δ*lafA* cell, shown in Fig. 4(h), presents segments in which the cell swims parallel and very close to the walls, followed by short segments in which the bacterium crosses from one wall to the opposite one. Most CHDs in this device occur when the bacteria either encounter a wall, or while swimming aligned to a wall. A minority of CHDs are detected when the bacterium is within one channel, away from the borders.

The distribution of reorientation angle Φ, in the inlets and in the artificial soil devices, for both strains are shown in Fig. 5(a). In the inlets, for both strains, the probability for a CHD to have an angle Φ ∈ [30°, 180°] is quite uniform, with only a small increase of approximately 5 % in probability around Φ = 40° and Φ = 180°, as compared to the probability at the center of the distribution. Note that small angles Φ *<* 30° are difficult to detect, so their probability is underestimated. The distribution does not change in the coarser soil with *w*_1_ = 20 μm. In the device with *w*_2_ = 10 μm, the probability distribution for the WT remains similar, but in the case of the mutant, there is an increase of probability for CHDs with large angle, Φ ≈ 180°. The trend is confirmed in the channels of width *w*_3_ = 5 μm, where there is an enlarged probability for Φ *>* 160° in both strains. This is consistent with the observed trajectories in this device, where RRs were common for bacteria swimming close to solid walls.

**Fig. 5.**
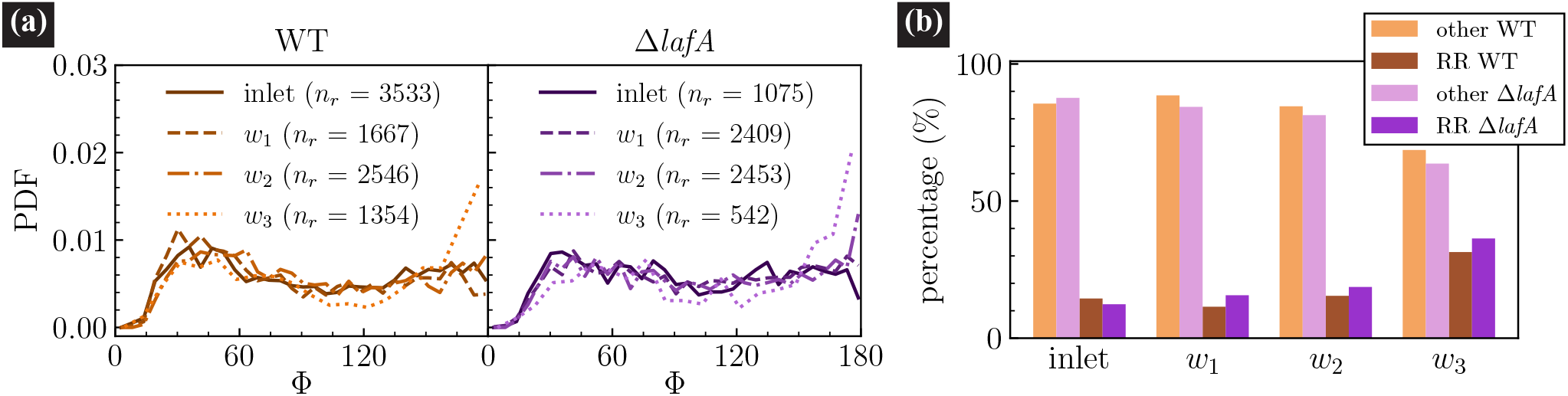
Angles and types of CHDs of *B. diazoefficiens* measured in the artificial soils. (a) Angle PDF of WT (left) and Δ*lafA* (right) in the inoculation chambers and in the different confined networks. (b) Classification of CHDs in the inlets and in the different soils, according to their angle: RR for Φ *>* 160°, and “other” otherwise.

Based on the threshold value stated above, we classified CHDs with Φ *>* 160° as RR, and the rest as “other”. As shown in Fig. 5(b), RRs amount to, approximately, 15 % of all CHDs for both strains in the inlets. The percentage of RR changes is marginal in the two widest artificial soils, but interestingly, it shows a significant increase in the soil with narrowest channels. There, approximately, 32 % of change of directions for the WT and 37 % for the single-flagellated Δ*lafA* correspond to Run and Reverses.

## Simulations of B. diazoefficiens in the soil

Thanks to a multidisciplinary work and to the detailed motility parameters measured exhaustively for both strains of *B. diazoefficiens*, their motion in confined geometries was simulated. Calculations were performed as realistically as possible, imitating both the exact microdevices design and the characteristics of their biological motors that propel them. The results described in the previous section were a part of our main original aims: to achieve a good statistical analysis, not found in the literature according to our knowledge, which is a key to success in the theoretical predictions using minimal models (see our previous Refs. Berdakin *et*.*al* (22; 23), and Montagna *et*.*al* (24)). The performed numerical research is done not only to complement and reproduce the experimental observation, but planning to go further and reach length and time scales that are unavailable to experiments in microfluidic devices and are essential for real soils and applications in agriculture.

We show in Fig. 6 examples of the arrays of grains used in the numerical calculations. Planar geometries exactly analogous to the experimental designs are used and shown in panel (a). In simulations the global size of the soil can be varied to perform scaling studies, but according to the experiments, the simulation area was kept with an aspect ratio of 2:1 (total length: total width). The channels width surrounding the impenetrable islands are fixed in each simulation, with width ranging between *w* = 5 μm and *w* = 20 μm as shown in the second row of Fig. 6(a).

**Fig. 6.**
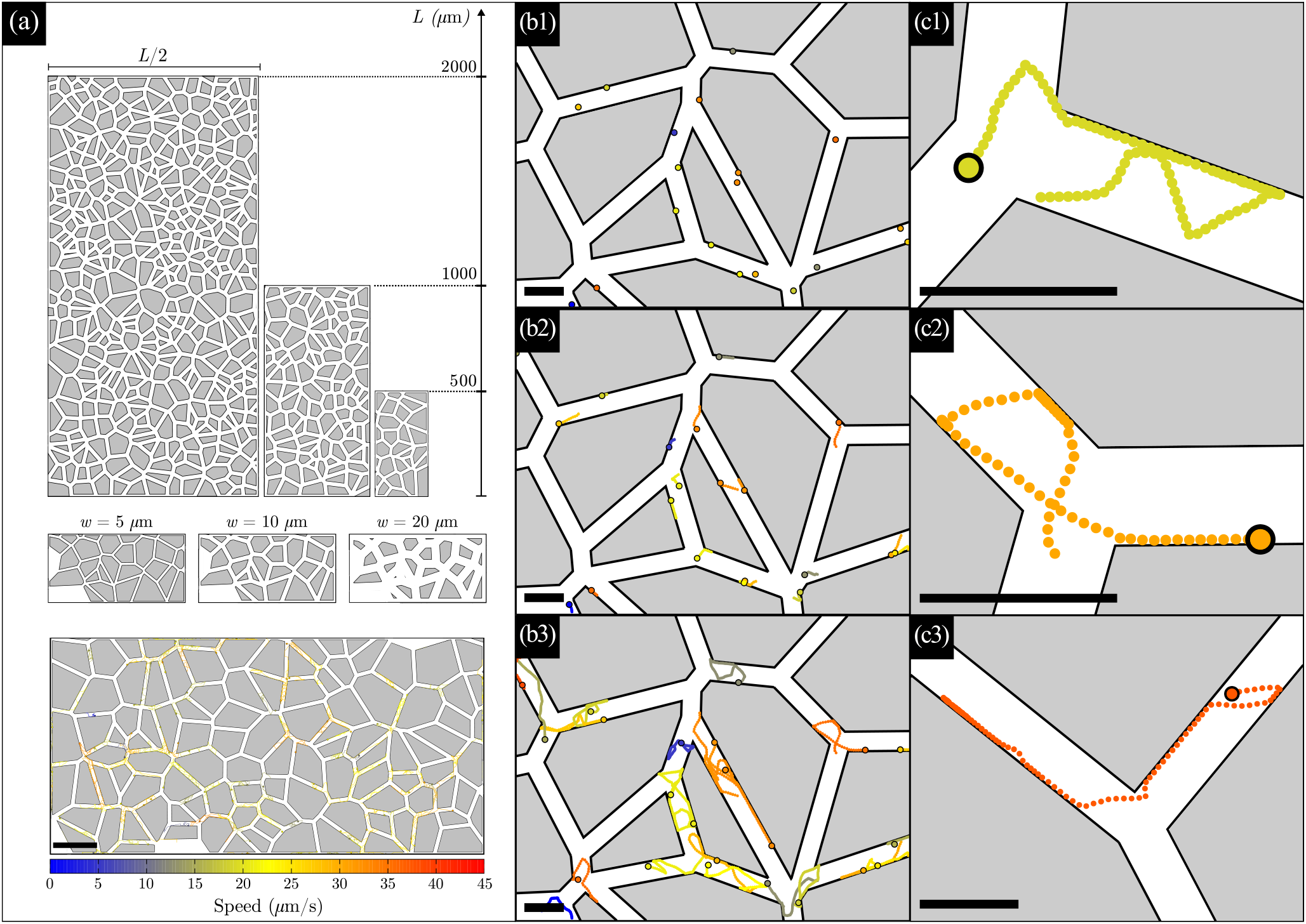
*B. diazoefficiens* swimming behavior in an *in-silico* soil-on-a-chip mimicking a porous media with 50 μm mean size grain. (a) Simulated arrays composed of islands and channels, with total area *L/*2 *× L*. In the second row, an array with *L* = 500 μm and three channel widths is shown: *w* = 5, 10, 20 μm (left to right respectively). In the lower panel, a few bacteria trajectories are drawn, inside a complete array with *L* = 1000 μm, *w* = 10 μm (scale bar of 100 μm). The color of the tracks correspond to the speed of each simulated bacterium, according to the speed color scale at the bottom (the same color scale is used in all panels). (b1) Initial positions, at *t* = 0. Dots representing bacteria are enlarged for visualization. (b2) Starting bacteria trajectories until *t* = 0.5 s. The black circles represent the final positions. (b3) Simulated tracks of duration 3.5 s. Most of the bacteria have performed several changes of direction, into the middle of the channels, or due to interactions. Note the swimming along walls, with some straight paths of more than 30 μm. (c1) to (c3) show three examples of simulated tracks, of duration 5.5 s, 5 s, and 3.75 s, respectively. Scale bars in (b) and (c) panels correspond to 20 μm.

Bacteria are located randomly inside the channels to start the simulations with distribution of speeds that imitates the measured speeds, Fig. 2(b)–(d),(f)–(h). A complete track of them into the simulated soil is shown in the lower part of Fig. 6(a), with the mean speed of each bacterium represented in colour. To improve visualization, only a fraction of the simulated bacteria are drawn. A zoomed-in region is shown in Fig. 6(b1-b3). In Fig. 6(b1) the initial positions are sketched and in Fig. 6(b2) the trajectories during the first steps simulated are shown. Specifically, at a time *t* = 0.5 s, most of simulated bacteria have not performed any CHD yet, and mostly straight paths are observed, of length proportional to the bacterial speed. In Fig. 6(b3) longer tracks, of more than three seconds, are presented in order to visualize more details. As observed experimentally, several CHDs and interactions with the walls are observed. In contrast to experimental tracks, numerical simulations have the advantage of not loosing any details close to walls or due to the swimming out of the focal plane. The CHDs are seen both close to boundaries, in the middle of the channels, or at the channel crossings. In addition, long tracks along walls can be detected with mean velocities of 30 μm*/*s for more than 30 μm.

Zooming in even more, several specifics tracks are presented in Fig. 6(c1) to (c3). Figure 6(c1) presents a single cell trajectory of total duration 5.5 s, where a clear RR of 180° occurs along the wall. In Fig. 6(c2), the bacterium moves from wall to wall, following a track, at first quite circular and later crossing more straight. Finally, Fig. 6(c3) shows a single long track that lasts 3.75 s. Note the longer track in shorter time due to the its larger velocity in comparison to the previous two examples. A long path following almost a full side of the grain is observed at the starting point, with a short detaching from the wall to come back to follow the wall. Later, a small CHD produces a channel crossing. At the end of the detection a large CHD is shown, very close to a RR. In brief, our model and simulations are able to reproduce the wide variety and complexity of the swimming behavior, in good agreement with the observed experimentally under confinement, as the examples shown in Fig. 4.

## Conclusions and perspectives

In this work, we present the measurement, with high detail, of the motility parameters of two strains of *B. diazoefficiens* in microfluidic devices that resemble the intricate geometry of porous soil. Our findings demonstrate that confinement reduces the average swimming speed of these bacteria (Fig. 2), a result that is likely caused by friction with solid walls and the increased interactions between bacteria and boundaries when confinement is important and comparable to their body sizes. These interactions also increase the fraction of run-and-reverse events in the distribution of changes of direction of bacteria (Fig. 5).

One of the main open questions regarding *B. diazoefficiens* swimming behavior concerns the function of its two flagellar systems. Consistent with previous studies (25), our observations indicate that, in unconfined situations, lateral flagella increase the swimming speed of the WT strain as compared to the Δ*lafA* mutant that lacks lateral flagella and only possesses the subpolar one. Additionally, our measurements in the inlets demonstrate that a larger speed diversity exists in the population of WT, which can be attributed to varying number and length of lateral flagella among individuals (Fig. 3(a)). Intriguingly, we observe that this larger diversity and higher speed of WT in comparison with the mutant disappear as microconfinement increases to 5 μm (Fig. 3(d)).

In synergy with the experimental findings, we developed a phenomenological numerical model. The simulations are able to reproduce with good agreement, Fig. 7, the dynamical behavior of a large population of micro-confined soil bacteria thanks to the use of realistic, experimentally measured parameters (Fig. 3(d)). This complement offers the opportunity to obtain further accurate predictions of transport properties in much larger length and time scales than those attainable with microfluidic devices, something that is fundamental to simulate real soils and field conditions (work currently in progress). Moreover, multiple spin-offs applied to other widely used inoculants of legume seeds are visualized, as well as the study of how bacterial motility is affected by different environmental factors.

**Fig. 7.**
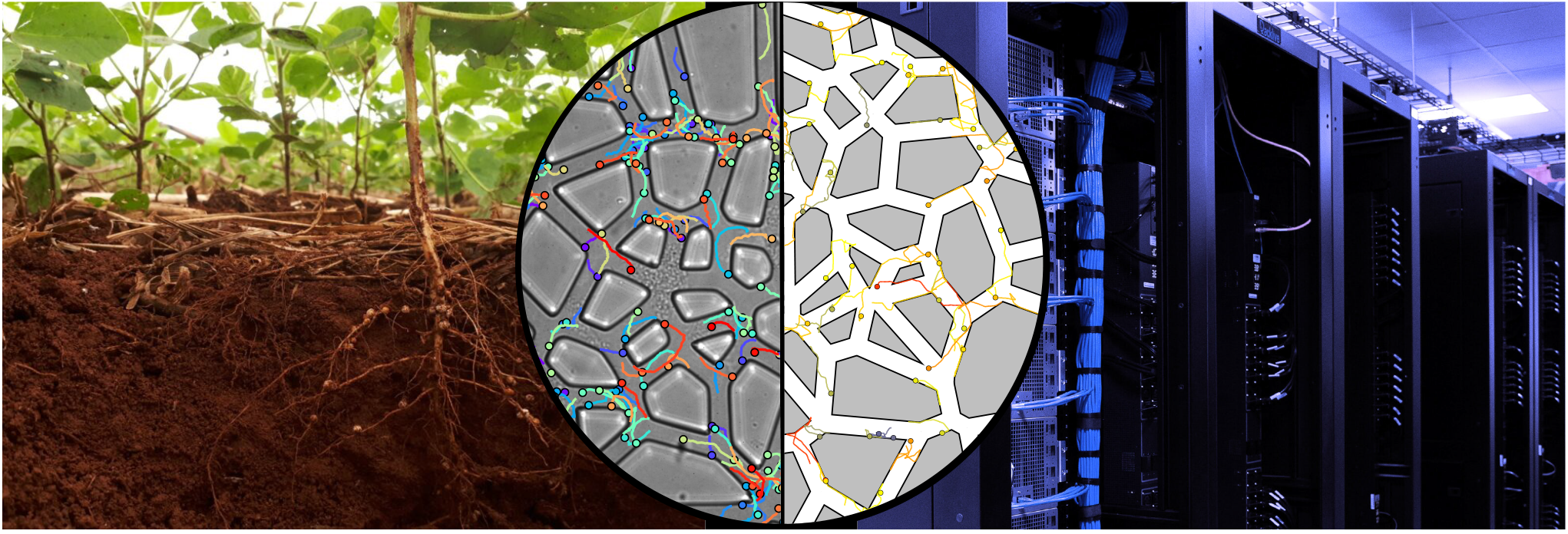
Soil-on-a-Chip for Studying *B. diazoefficiens* Confined Swimming behavior: Experimental vs Theoretical achievements. The central circle highlights the reached goals of this work: good agreements were found between real-time microbial tracking in microfludic devices and predictive theoretical models and calculations, utilizing high-performance computing (HPC).

Biological nitrogen fixation, such as the one performed in the symbiosis between *B. diazoefficiens* and soybean, can play a fundamental role in the development of greener crops by substituting artificial fertilizers that destabilize Nitrogen balance in the soil (17). Sustainable intensive agriculture is indeed required to face the challenges of food and environmental safety of the future generations, and for that, social, political and technological innovations are required (26; 16). Bioproducts as rhizobia inoculants have been in the market for decades now, but are still far from competitive in comparison with contaminant but more efficient chemical products. Our work combines multidisciplinary contributions from microfabrication technologies, fluids and computational physics and genetic-agronomic background to better understand the behavior of *B. diazoefficiens* in the soil and contribute to the development of a new generation of biofertilizers. So far, the motility performance of different strains could only be compared directly by microscopy in simple homogeneous liquid medium, or by measuring colony growth on agar plates, where bacteria move by swarming instead of swimming. Valuable but indirect information can also be obtained by recovering bacteria in roots or soil samples. Our biocompatible and transparent microfluidic soil-on-a-chip devices, on the contrary, allow to directly visualize bacteria behavior in soil-like conditions, providing valuable insights into the behavior of this beneficial bacterium in controlled and replicable experiments and calculations. Even more, our findings are statistically detailed and constitute a clear evidence of the power that microfludics SOCs tools and multidisciplinary approaches, Fig. 7, can offer to the development of sustainable agriculture practices.

## Methods

### Microfludic devices

Microfluidic devices were fabricated using standard lithography and soft lithography methods. Templates were fabricated using optical lithography on 2” silicon wafers using SU-8 (Gersteltec Sarl) with a maksless laser writer (MLA100, Heidelberg Instruments). From the molds, polydimethylsiloxane (PDMS, Sylgard 184, Dow Corning) replicas were obtained. Once cured, the PDMS device was cut with a scalpel and carefully detached from the molds. Inlet and outlet holes were pierced in the PDMS blocks with a 1.5 mm-diameter biopsy punch, after which both the PDMS block and a microscope glass slide were exposed to an air plasma for one minute and sealed together.

The devices consisted of an inoculation chamber, a region with a network of narrow, randomly oriented channels, and an exit chamber. The network was designed in Matlab as a Voronoi tessellation, obtained from *N* = 3000 seed points within a region of *A* = 2 × 1 (arbitrary units) (27; 28). To generate a disordered network, these points were required to be at least 0.3*d*_0_ units apart, where 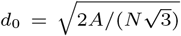 would be the distance between the seed points in a completely regular network (a honeycomb network), but other than that they were randomly generated. The edges of the Voronoi polygons were then given a fixed width and the network was re-scaled to real dimensions. The same design was used for the three networks, which differed in their scaling factor and final height (defined in the lithographic step). The final dimensions (height *h*_*i*_, channel width *w*_*i*_ and overall area of the network *L*_*i*_ × *W*_*i*_, *i* = 1, 2, 3) of the devices were (i) *h*_1_ = 25 μm, *w*_1_ = 20 μm and area 4 × 2 mm^2^, (ii) *h*_2_ = 25 μm, *w*_2_ = 10 μm and area 2 × 1 mm^2^, and (iii) *h*_3_ = 15 μm, *w*_3_ = 5 μm and area 1 × 0.5 mm^2^.

### Bacterial strains and culture

*B. diazoefficiens* USDA 110 (obtained from the United States Department of Agriculture, Beltsville) is the wild type strain of the species. The derived mutants Δ*fliC* and Δ*lafA*, which are devoid of subpolar or lateral flagella respectively, were described elsewhere (29).

Bacteria were grown in HM salts medium (30), supplemented with 1 g*/*L yeast extract and 5 g*/*L L-arabinose (HMY-Ara), as described (10). For routine use, bacteria stocks where kept at 4 °C in yeast extract mannitol-agar (1.5 %) (31), supplemented with chloramphenicol 20 mg*/*L plus kanamycin 200 mg*/*L (Δ*fliC*) or streptomycin 400 mg*/*L (Δ*lafA*). Cultures in agar plates were renewed every 3 months from frozen stocks kept at −80 °C in culture medium HMY-Ara supplemented with 20% v/v glycerol. Liquid cultures were initiated from a single colony in HMY-Ara and grown for seven days at 28 °C and 180 rpm until OD_600_ = 3.0 ± 0.1. A 1:100 dilution into HMY-Ara was then grown for two days at the same temperature and agitation conditions until OD_600_ = 1 ± 0.1. A 1:100 dilution into HM supplemented with L-arabinose 5 g*/*L (HM-Ara) was then left at 28° without agitation for a minimum of 6 h before chamber inoculation.

At least three different cultures for each strain starting from different colonies in different days (biological replicas) were used to ensure reproducibility of the experiments.

### Inoculation and data acquisition

Microfluidic devices were filled with HM-Ara medium prior to use in order to eliminate air bubbles. Bacteria were inoculated into the devices at constant flow rates, between 1.6 nL*/*s and 30 nL*/*s, using 1 mL glass syringes (Hamilton Co.), a syringe pump (neMESYS Base 120 and low pressure module V2, Cetoni) and Tygon tubes (0.02” ID x 0.06” OD Microbore Transfer Tubing, Masterflex) until bacteria had completely entered the device. Tubing were then cut near the syringe needle and the tubing ends were introduced in a reservoir with HM-Ara. After an equilibration time, required to relax any remaining flow, bacteria were observed in bright field using an inverted microscope (TS100 LED, Nikon) equipped with a 40X microscope objective and a CMOS camera (Zyla 4.2, Andor or Mini AX100, Photron). The spatial resolution of the optical system was 6.24 pixel*/*μm with the Andor camera and 3.06 pixel*/*μm with the Photron camera. The effective depth of field was determined to be 4 μm by fixing the focus at bacteria attached to the glass surface and slowly moving the objective until the attached bacteria were no longer detectable. It is important to remark that the focus depth is merely four times the bacteria dimensions, then they can cross it just in 0.18 seconds if they swim at 22 μm*/*s. Several 60 s image sequences were recorded at 50 fps, for later analysis, focusing at different sites of the network but also at the inoculation and exit chambers.

In principle, from a given culture, three aliquots were used to inoculate the three networks during the same day. However, when statistics were not deemed enough for a given network and/or strain, additional measurements were obtained with a different culture in a specific network.

### Bacteria tracking

The bacteria trajectories along time were obtained through image analysis of the experimental videos. The positions were extracted using the open software *Biotracker*, developed within our group of collaborators in FaMAF-National University of Córdoba (32; 33). BioTracker is an open-source computer vision framework designed for visual cell tracking. The software is suitable for a variety of image recording conditions and offers a number of different tracking algorithms.

Several well-known particle tracking software available are built generically, and may not be optimized for fast microswimmers or small cells that are hard to detect. To address this issue we developed BioTracker specifically to follow fast and sub-micron sized microswimmers, with specific motility features such as changes of direction along their paths or complex 3D oscillations of sperm cells (34). In 2016, when the software was developed, tracking fast-moving bacteria with high precision, without confusing the tracks’ projections and crossings, was a significant challenge. Despite this challenge, the software was able to deliver better results than 14 methods compared in the article by Chenouard *et. al*, published in Nature Methods 2014 (35). The software utilizes Kalman Filter estimators in its linker module, which contributed to its success in tracking fast-moving bacteria.

The BioTracker software offers several advantages in analyzing with precision videos of long duration. It demonstrates stability in performance over various metrics and scenarios, low computational cost, and robustness. Specifically built to track microswimmers with rapid and unpredictable movements, including fast and abrupt runs and tumbles, BioTracker *automatically counts and characterizes such events and their angles* in a dedicated module. Its success is attributed to its detector module and the qualified post-acquisition analysis of subtle details of the tracks, such as small changes of directions or run and reverse events, which are extensively used in the research presented here. The software provides a user-friendly interface and supports the implementation of new problem-specific tracking algorithms, allowing the scientific community to accelerate their research and focus on the development of new vision algorithms.

A change of direction is detected in a given position if there is a change of the swimming orientation with respect to the swimming orientation a number of bodies (*n*_*b*_) behind and ahead from the position, where the number *n*_*b*_ is chosen by the user. For a correct implementation, it is fundamental to have a good characterization of the cell body sizes. Moreover, the population can be heterogeneous, and BioTracker takes that into account. A lower threshold for the CHD angle was defined at 15°, to distinguish from usual Brownian motion.

In this work, we were able to detect and track thousands of soil diminutive bacteria simultaneously and precisely extract all their motility parameters, not just their main speed, using BioTracker. By fine-tuning the software options, we were able to achieve precise confined motility measurements of *Bradyrizobium*, a bacterium that is extremely small and hard to detect and follow in-plane, and even harder to detect their Run-and-Reverses, events that occur very fast in distances of the order of their body sizes. The results showed in previous sections demonstrate the advantages of BioTracker (32; 33) and the successful final achievements in tracking and characterizing in SOCs the motility of these soil microbials.

### Model and simulations

We simulate the behavior of soil bacteria diluted in an aqueous medium and confined to artificial soils at low Reynolds numbers (36). This system is characterized predominantly by viscous effects acting on cells. We model *N*_*b*_ bacteria, reproducing the concentration of cells inoculated in each experiment, and assume they are self-propelled thanks to specific motor forces proportional to their speeds, *v*. Each strain has its unique swimming strategy, which we incorporate into the model. Our aim is to develop a simple yet extremely accurate model that can efficiently simulate large populations of *N*_*b*_ microswimmers. As bacteria swim inside SOCs, surfaces dominate, and interactions with walls significantly impact the simulation’s CPU time. To achieve realism, we utilize measured data for the motility parameters of each strain (Figs. 3 and 5) without approximating biological parameters in the calculations.

The phenomenological model we propose for the confined bacteria behavior shares similarities with previous successful proposals for other bacteria swimming in microdevices or porous media (22; 23; 24). The dominant forces we consider are: 1. Motor force, **F**^*m*^. 2. Bacteria interactions with walls, **F**^*bw*^. 3. Fluid damping force proportional to speed and −*γ*, the damping constant in an aqueous medium. Although the system is diluted, we also account for soft interactions among bacteria, **F**^*bb*^, for completeness.

Here, we adapted and optimized previous models to reduce computing time, accommodating the geometry of SOCs and the speed and specific change of directions distributions of *B. diazoefficiens*, as Run-and-Reverse movements, which are crucial. The initial positions are randomly distributed across the device (see Fig. 6), while the initial speeds follow the same measured speed distributions (Fig. 3). From this starting point, we have a predictive set of 2*N*_*b*_ equations of motion. The over-damped motion equation for each soil bacterium, positioned at **r** and resulting from the cancellation of the sum of forces acting on its body, is as follows: *γd***r***/dt* = **F**^*m*^ + **F**^*bw*^ + **F**^*bb*^. In addition, a coupled time-evolution equation for the velocity direction, is added, where the measured change of directions are imposed (22; 23; 24).

The bacteria are modeled as disks with a radius of *R*_*b*_ = 0.5 μm, because the averaged measured diameter is one micron. We use real measured parameters to propose the equations of motion for positions and velocity direction, as described in Refs. (22; 23; 24). The equations of motion are solved using Langevin dynamics methods. We introduce the exact change of direction distribution measured, as shown in Fig. 5, using the rejection method of Von Neumann to reproduce the distributions exactly, rather than relying on approximations commonly used in the literature and our previous contributions. We also optimize the codes to speed up calculations of bacteria-grain interactions, resulting in a 50% improvement in performance compared to previous codes (optimization modules are available; contact the corresponding authors). As a result, the confined trajectories at all times are obtained in our variety of soil-on-a-chip devices with different geometries and channels widths (Fig. 6), in a good experimental-numerical agreement (Fig. 7).

## Acknowledgements

This work was partially funded by Secyt-UNC, CONICET, PICT 2015-0735, PICT-2020-SERIEA-0293 (Argentina), and ANID - Millennium Science Initiative Program - NCN19_170 (Chile). Fabrication of microfluidic devices was possible thanks to ANID Fondequip grants EQM140055 and EQM180009. Simulations were done thanks to GTMC-group computational facilities in FAMAF-UNC, Argentina. MPM Acknowledges funding from Fondecyt postdoctoral grant No. 3190637. Experimental assistance from Francisca Paredes Oltra and a careful reading of the manuscript from Pedro Pury and Adolfo J. Banchio is greatly appreciated, as well as useful discussions.

## Author contributions statement

VIM, MLC, and AL conceived the work. MPM and JPC conducted the experiments. NG, MPM, JPC, and SM analyzed experimental data. VIM developed the model and offered the Biotracker. NG, SM, and VIM performed simulations. MLC, VIM and AL wrote the manuscript. All authors reviewed the manuscript.

